# Methionine synthase is essential for cancer cell proliferation in physiological folate environments

**DOI:** 10.1101/2020.06.12.149005

**Authors:** Mark R. Sullivan, Alicia M. Darnell, Montana F. Reilly, Caroline A. Lewis, Matthew G. Vander Heiden

## Abstract

Targeting folate metabolism can be an effective way to treat cancer. The enzyme methionine synthase catalyzes a key reaction in both folate and methionine metabolism. Early work suggested that inhibiting methionine synthase might restrain tumor growth, though the mechanism remains unclear. We find that due to its specific role in processing folates, methionine synthase is required for cancer proliferation. However, widely used cell culture conditions obscure the proliferative and metabolic consequences of methionine synthase inhibition. Complete dependence on methionine synthase only arises when 5-methyl tetrahydrofolate, the major folate found in circulation, is the predominant folate source provided to cells. In these physiological folate conditions, methionine synthase activity is necessary to maintain intracellular levels of nucleotides, but not methionine. These data reveal that the extracellular environment can alter the essentiality of methionine synthase and suggest that this enzyme plays a crucial cell-autonomous role in supporting nucleotide synthesis and cell proliferation in physiological contexts.

## INTRODUCTION

Folate metabolism is critical for cell proliferation and survival. One-carbon units carried by folate cofactors are necessary for many one-carbon transfer reactions^1^, including those required for nucleotide synthesis^2^. Folate species carry distinct one-carbon units in different oxidation states to support these biosynthetic reactions, with each folate species derived in cells from a common tetrahydrofolate (THF) scaffold (Figure 1A). THF cannot be synthesized by human cells de novo^3^; thus, mammalian cells rely on uptake of exogenous precursors which can be used to generate THF and enable the one-carbon transfer reactions that support proliferation. The available THF precursor differs between standard cell culture media and what is found in circulation in mammals. In standard culture, cells are provided with folic acid. In mammals, the predominant circulating folate is 5-methyl tetrahydrofolate (5-methyl THF)^3,4^. As environmental nutrients can strongly influence cellular metabolism^5,6^, we sought to understand how culturing cells in 5-methyl THF, the physiological folate source, would impact intracellular metabolite levels and dependence on metabolic pathways that utilize folate species.

**Figure 1.**
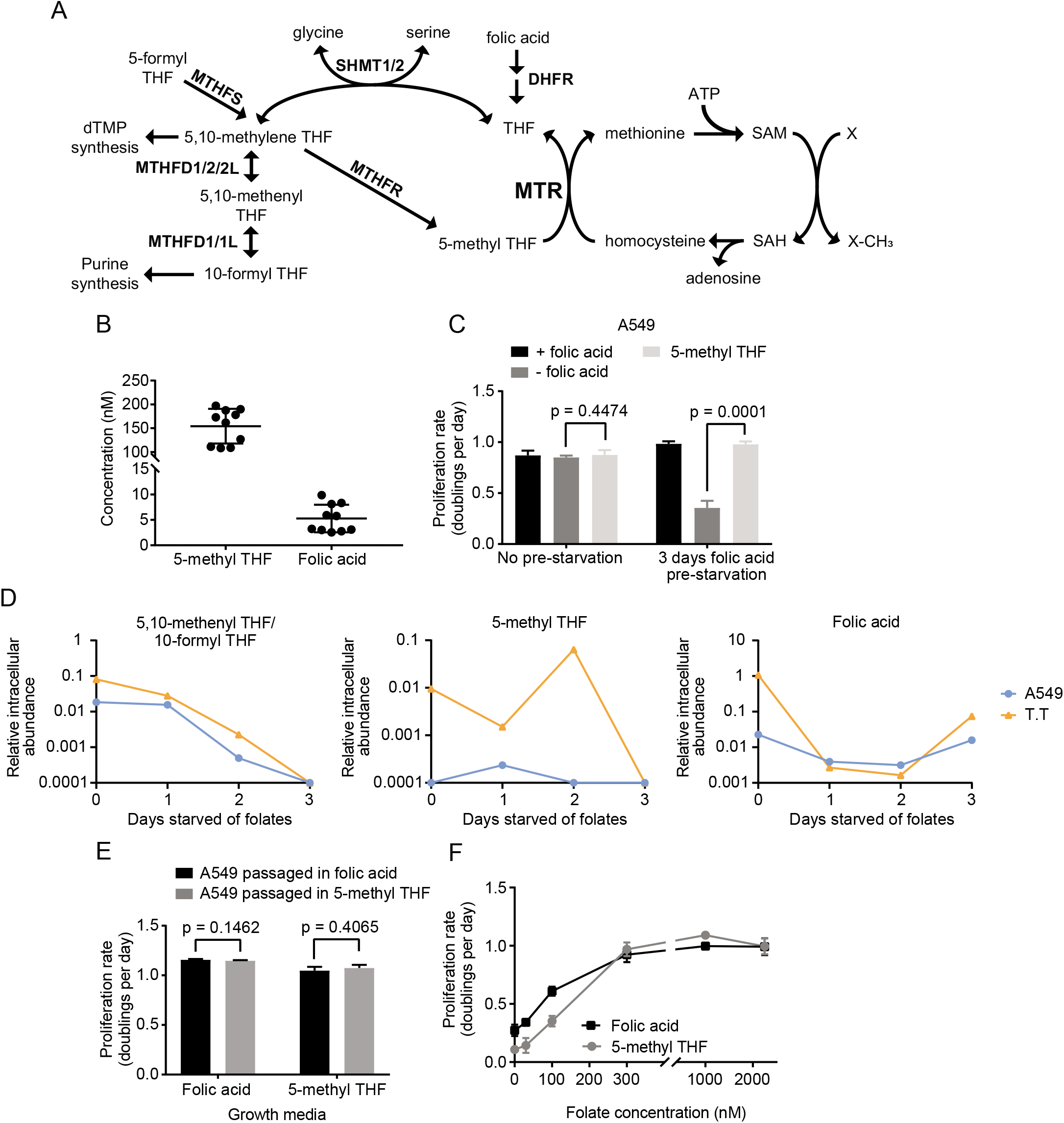
Cells can be cultured with 5-methyl THF as the only folate source. **(A)** Schematic showing reactions involved in folate and methionine metabolism. SAM: S-adenosyl methionine. SAH: S-adenosyl homocysteine. MTR: methionine synthase. DHFR: dihydrofolate reductase. SHMT: serine hydroxymethyl transferase. MTHFR: methylenetetrahydrofolate reductase. MTHFD: methylenetetrahydrofolate dehydrogenase. MTHFS: methylenetetrahydrofolate synthetase. dTMP: deoxythymidine monophosphate. **(B)** LC/MS measurement of 5-methyl THF and folic acid levels in plasma from NSG mice. n = 10 mice. **(C)** Proliferation rate of A549 cells in media with the indicated folate before (No pre-starvation) or after 3 days of culture with no folic acid (3 days folic acid pre-starvation). **(D)** LC/MS measurement of intracellular 5,10-methenyl THF/10-formyl THF, 5-methyl THF, and folic acid levels in A549 or T.T cells cultured for the indicated number of days in media lacking any added folates. Data are normalized to cell number and an internal standard. **(E)** Proliferation rate of A549 cells when switched to media containing folic acid or 5-methyl THF after 3 weeks of continuous culture in media containing either folic acid or 5-methyl THF as the only folate as indicated. **(F)** Proliferation rate of A549 cells in media containing the indicated amounts of either folic acid or 5-methyl THF. Mean +/- SD is displayed for all panels. p values indicated on all panels are derived from two-tailed, unpaired Welch’s t tests.

Cells use different pathways to convert folic acid and 5-methyl THF to THF for use in subsequent biosynthetic reactions^3^ (Figure 1A). Folic acid is directly converted to THF by the enzyme dihydrofolate reductase (DHFR). In contrast, the conversion of 5-methyl THF to THF is coupled to synthesis of the amino acid methionine by transfer of a methyl group to homocysteine in a reaction catalyzed by the vitamin B12 (cobalamin)-dependent enzyme methionine synthase (MTR)^7,8^ (Figure 1A). MTR is the only known enzyme that can utilize 5-methyl THF as a substrate and regenerate the THF scaffold. Therefore, loss of MTR activity secondary to cobalamin deficiency causes accumulation of intracellular 5-methyl THF, clinically referred to as the “methyl-folate trap”^9,10^. This irreversible sequestration of folates in the form of 5-methyl THF in cells results in limited THF availability to accept new one-carbon units. Notably, culturing cells in folic acid rather than 5-methyl THF bypasses the requirement for MTR to regenerate THF, and may therefore mask dependence on MTR to support proliferation.

Several lines of evidence suggest that MTR may be critical for tumor proliferation. Loss of MTR activity due to impairment of cobalamin binding or availability perturbs the folate cycle and slows cancer cell proliferation^11,12^. Further, clues from human medicine suggest that cobalamin metabolism may be important to sustain tumor growth^13–15^; however, cobalamin acts as a cofactor for other metabolic pathways^16^, which confounds interpretation of these results. MTR activity also impacts processes beyond the folate cycle, including regeneration of the methyl donor cofactor S-adenosyl methionine (SAM)^9,17–19^ (Figure 1A). In animals, MTR is important for maintaining whole-body methionine and SAM levels^9,20–22^, and MTR inhibition has also been suggested to impair cancer cell proliferation in cultured cells by specifically interfering with production of these metabolites^17,18^. However, whether targeted MTR inhibition would curtail tumor growth in animals is unclear, and the metabolic consequences of perturbing MTR activity have not been examined under physiological folate conditions.

## RESULTS

To understand the role of MTR in folate metabolism, methionine metabolism, and tumor growth, we examined the metabolic and proliferative consequences of MTR deletion in physiological folate contexts *in vitro* and *in vivo*. The predominant folate in circulation in humans is 5-methyl THF^4^, and we confirmed that this is also true in mice fed a standard chow diet (Figure 1B). To accurately model folate metabolism in cultured cells, we cultured human cancer cells in media where 5-methyl THF is the sole folate source and characterized the effect of different folate sources on cell proliferation. Previous work has shown that leukemia cells can be cultured for short periods in 5-methyl THF^11,17,18^. To generalize these findings to cancer cells derived from solid tumors, we examined A549 and T.T cells, which exhibit average MTR expression levels ^23^ (Supplemental Figure 1A). We observed that folate deprivation did not diminish proliferation of either cell line over four days of exponential growth in culture. (Figure 1C, Supplemental Figure 1B). This suggests that cells contain excess folates that are sufficient to sustain proliferation over this period and is consistent with previous observations in leukemia cells^11^. Additionally, the dialyzed fetal bovine serum used to supplement the culture media contains low levels of both folic acid and 5-methyl THF (Supplemental Figure 1C), providing a potential alternative source of some folates. In order to generate a system to study cells in conditions with well-defined sources of folates, we pre-incubated cells in media lacking any added folic acid beyond what is found in dialyzed serum in an attempt to deplete intracellular folate reserves before assessing cell proliferation and metabolism. Three days of such folate deprivation was sufficient to deplete intracellular levels of the folate species 5,10-methenyl THF/10-formyl THF and 5-methyl THF (Figure 1D) without reducing the levels of intracellular amino acids, SAM, or nucleotides (Supplemental Figure 1D-F) or impairing proliferation rate (Supplemental Figure 1G). This suggests that the cells have consumed nearly all of their intracellular folate stores over three days of culture in folate-depleted media, but that their metabolism remains largely unaffected. Thus, we reasoned that short-term folate deprivation would allow for controlled replacement of the environmental folate source.

To examine the metabolism of cells that have access to 5-methyl THF as their only folate source, cells were cultured for three days in folate depleted media and then cultured for four additional days in media with either folic acid, 5-methyl THF, or no folates supplemented. Cells are able to proliferate at the same rate when cultured in media with either folic acid or 5-methyl THF as the folate source, but cells deprived of folates for this extended period of time display a proliferation defect (Figure 1C, Supplemental Figure 1B). Importantly, cells maintain the same proliferation rate even after culture in 5-methyl THF for 3 weeks, and this proliferation rate remains unchanged when cells are switched back to standard media containing folic acid as the folate source (Figure 1E). These data confirm that providing 5-methyl THF as the sole folate source does not alter the long-term proliferative capacity of the epithelial cancer cells considered in this study. Further, 5-methyl THF and folic acid sustain proliferation at similar rates when exposed to different concentrations of either folate source (Figure 1F). Of note, the average concentration of 5-methyl THF in human plasma is 37.5 +/- 1.5 nM^4^ and in mouse plasma is 154.2 +/- 26.1 nM (Figure 1B), neither of which is able to sustain maximal proliferation of A549 cancer cells in culture (Figure 1F). This suggests that folate availability may limit rapid proliferation in some physiological settings, consistent with reports that folate supplementation can accelerate the growth of some cancers^24–26^. Together, these results suggest that 5-methyl THF is sufficient to sustain the growth of human cancer cells in the absence of other folate sources.

As cancer cells are able to proliferate in culture with 5-methyl THF as the sole folate source, we next used this system to examine the role of MTR when 5-methyl THF is the major folate source. For these studies, we disrupted MTR expression in A549 and T.T cells using CRISPR-Cas9 genome editing (sgMTR) and then re-expressed an sgRNA-resistant version of MTR (+MTR) or an empty-vector control (EV) to create isogenic cell lines that differ in MTR expression (Figure 2A). In both A549 and T.T cells, MTR is essential for proliferation when 5-methyl THF is the sole folate source (Figure 2B-C). MTR knockout cells display high steadystate levels of 5-methyl THF regardless of folate source (Figure 2D-E), consistent with previous results^27^. This trapping of folate as 5-methyl THF in the MTR knockout cells may explain the small decrease in proliferation observed in A549 MTR knockout cells compared to EV control cells when cultured in folic acid. As expected, levels of 5-methyl THF remained high in MTR knockout cells even upon folic acid starvation (Supplemental Figure 2A). Further, other intracellular folate species that are synthesized using THF were depleted more rapidly in MTR knockout cells than control cells following folate starvation. Consistent with this observation, MTR knockout cells were more sensitive to short term folate starvation than control cells (Supplemental Figure 2B). Thus, proliferation of MTR knockout cells can be limited by folate availability, and these cells do not require pre-starvation of folates before replacement of the environmental folate source. Interestingly, growth in 5-methyl THF also led to accumulation of 5-formyl THF (Figure 2D-E), a folate cofactor that may alter the activity of other folate-dependent enzymes^27^, suggesting that growth in 5-methyl THF may broadly affect intracellular folate metabolism.

**Figure 2.**
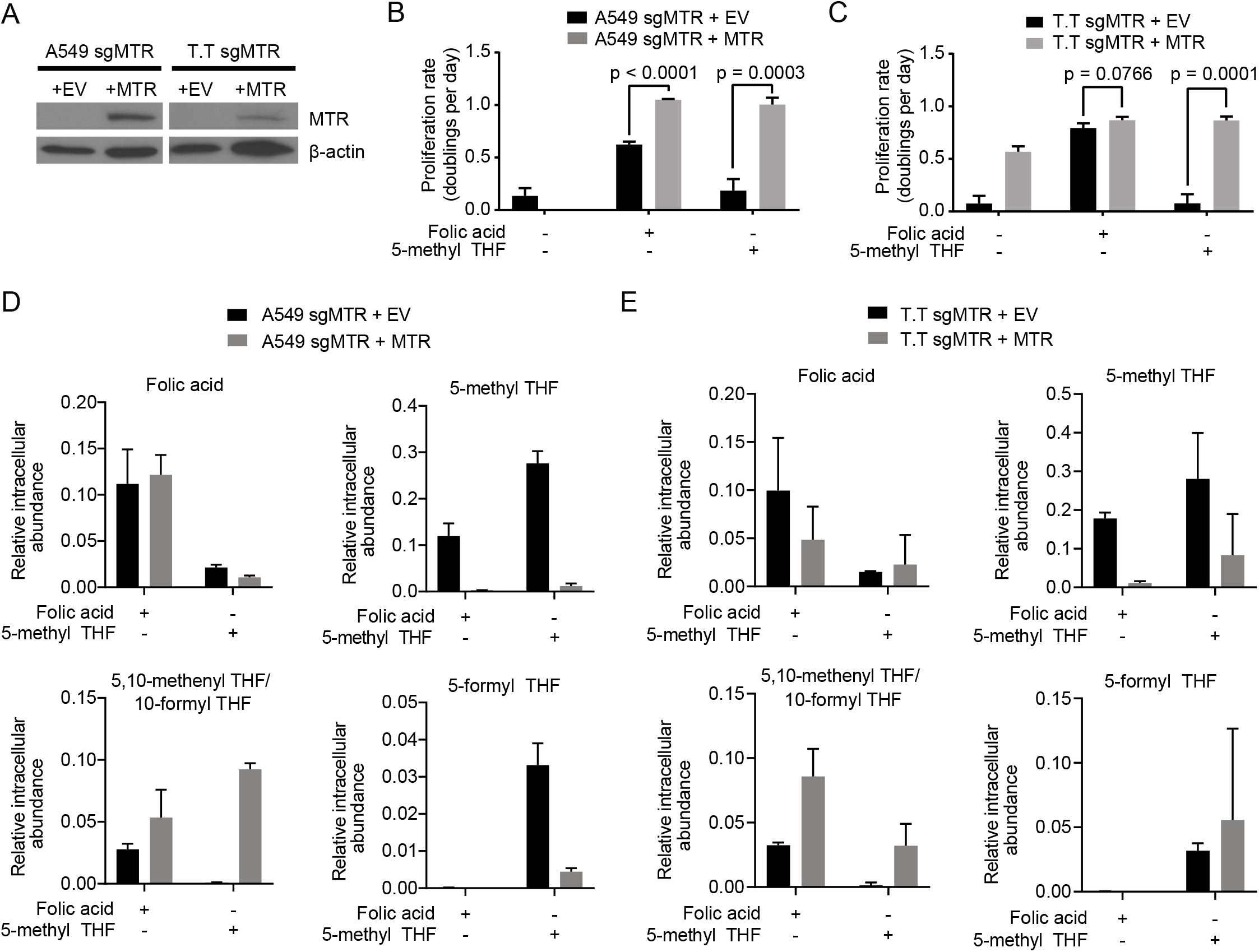
Methionine synthase is essential for proliferation if 5-methyl THF is the only folate source. **(A)** Western blot analysis of MTR expression in A549 and T.T cells targeted with an sgRNA to disrupt MTR expression and then engineered to either express an empty vector (+EV) or an sgRNA resistant MTR cDNA (+MTR). Proliferation rates of A549 **(B)** and T.T **(C)** cells described in A without (+EV) or with MTR expression (+MTR). Cells were cultured in the indicated folate for 4 days after 3 days of folate pre-starvation. **(D)** LC/MS measurement of intracellular folic acid, 5-methyl THF, 5,10-methenyl THF/10-formyl THF, and 5-formyl THF levels in A549 cells **(D)** or T.T cells **(E)** without (+EV) or with MTR expression (+MTR) cultured for four days in media with the indicated folate. LC/MS data are normalized to cell number and to an internal standard. Mean +/- SD is displayed for all panels. p values indicated on all panels are derived from two-tailed, unpaired Welch’s t tests.

To understand the role of MTR in supporting proliferation in 5-methyl THF-containing media, we examined the metabolic consequences of MTR loss. MTR can support both methionine and folate metabolism. If insufficient intracellular methionine levels upon MTR loss suppress proliferation, addition of exogenous methionine may rescue proliferation. The standard culture conditions used in this study contain 100 μM methionine, a concentration similar to that found in murine plasma^28^. Providing additional methionine does not rescue proliferation of MTR knockout cells (Figure 3A, Supplemental Figure 3A). Indeed, methionine levels are higher in MTR knockout cells than control cells (Figure 3B, Supplemental Figure 3B). This accumulation of methionine is likely a non-specific phenomenon, as nonessential amino acids such as serine and essential amino acids such as lysine also accumulate in these cells (Figure 3B, Supplemental Figure 3B). Together, these data argue that MTR knockout does not compromise intracellular methionine levels in physiological extracellular folate and methionine conditions.

**Figure 3.**
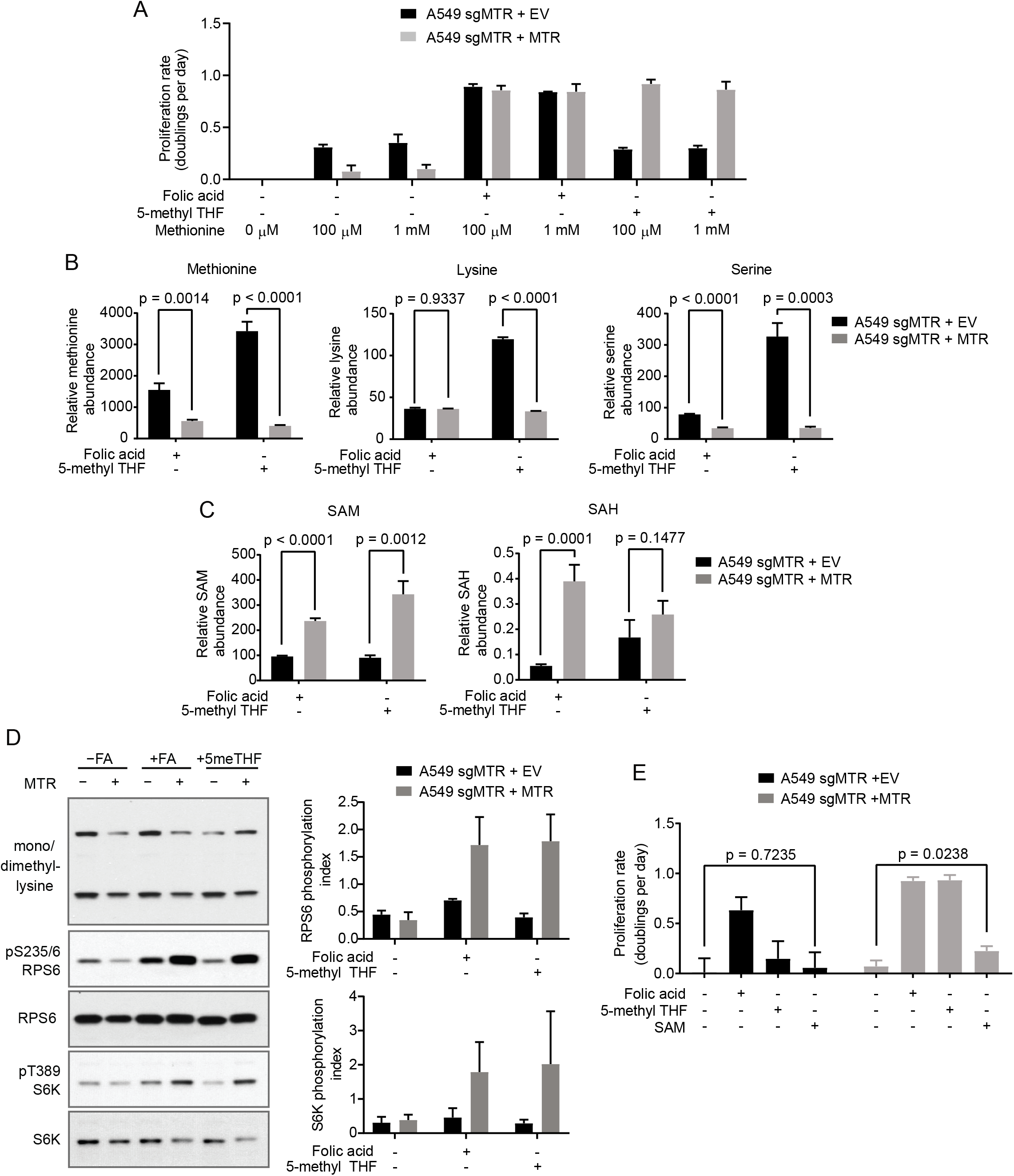
MTR knockout reduces SAM levels but not methionine levels. **(A)** Proliferation rates of A549 cells without (+EV) or with MTR expression (+MTR) cultured in the indicated folate with or without the addition of 1 mM methionine to supplement the 100 μM methionine present in RPMI-1640 culture media. **(B)** LC/MS measurement of intracellular methionine, lysine, and serine levels in A549 cells without (+EV) or with MTR expression (+MTR) cultured for 4 days in the indicated folate. Data are normalized to the total protein content of cells in each condition and to an internal standard. **(C)** LC/MS measurement of intracellular SAM and SAH levels in A549 cells without (+EV) or with MTR expression (+MTR) cultured for 4 days in the indicated folate. Data are normalized to the total protein content of cells in each condition and to an internal standard. **(D)** Western blots to assess levels of mono- or dimethyl lysine-containing proteins, RPS6 phosphorylated at S235/6, total RPS6, S6K phosphorylated at T389, and total S6K in A549 cells without (+EV) or with MTR expression (+MTR). Phosphorylation index is defined as the phosphorylated protein antibody signal divided by the same total protein antibody signal in each lane. **(E)** Proliferation rates of A549 cells without (+EV) or with MTR expression (+MTR) cultured in the indicated folate with or without the addition of 1 mM SAM. Mean +/- SD is displayed for all panels. p values indicated on all panels are derived from two-tailed, unpaired Welch’s t tests.

Methionine is directly converted to SAM by addition of adenosine triphosphate (ATP) (Figure 1A). In contrast to methionine, SAM levels and S-adenosyl homocysteine (SAH) levels are decreased in MTR knockout cells (Figure 3C, Supplemental Figure 3C) and could thus contribute to the proliferation defect observed in cells lacking MTR and cultured in 5-methyl THF. As SAM is used as a methyl donor for methylation reactions in cells, SAM levels could affect proliferation by altering protein or DNA methylation patterns^29,30^. SAM has also been shown to signal to mTORC1, and reduced SAM might affect proliferation by inhibiting mTORC1 kinase^31^. However, SAM levels are equally diminished in MTR knockout cells that are proliferating in folic acid and those cultured in 5-methyl THF (Figure 3C, Supplemental Figure 3C), arguing that SAM depletion does not explain why MTR knockout cells cannot proliferate in 5-methyl THF. Changes in methylation of cytosolic proteins did not correlate with the reduced SAM levels or proliferation rate caused by MTR knockout in 5-methyl THF (Figure 3D, Supplemental Figure 3D), suggesting that this is unlikely to be responsible for the proliferation defect in these cells. Phosphorylation of the direct mTORC1 kinase target S6 kinase, as well as phosphorylation of the S6 kinase target RPS6, largely correlated with SAM levels (Figure 3D, Supplemental Figure 3D). To determine whether SAM-mediated changes to mTORC1 signaling might explain the growth defect of MTR knockout cells in 5-methyl THF, we tested whether addition of exogenous SAM was sufficient to rescue growth. Providing exogenous SAM can reactivate mTORC1 signaling in cultured cells^31^ but is unable to restore MTR knockout cell proliferation in 5-methyl THF (Figure 3E, Supplemental Figure 3E). Together, these experiments suggest that MTR expression is important to maintain levels of SAM and SAH in cells, but that the decreases in SAM and SAH levels observed with MTR knockout are not sufficient to explain reduced cell proliferation in physiological folate conditions.

MTR is required to convert 5-methyl THF to THF, allowing THF to accept new one-carbon units and generate the folate species required for nucleotide synthesis. As such, MTR inactivation has been linked to impaired pyrimidine nucleotide synthesis^27^. Thus, we sought to examine the effects of MTR knockout on folate species required for nucleotide synthesis. The folate precursor for purine synthesis, 10-formyl THF, is indistinguishable from a closely related species 5,10-methenyl THF by LC/MS due to known interconversion after cell lysis^32,33^. We observed that MTR knockout results in depletion of the combined levels of 5,10-methenyl THF and 10-formyl THF (Figure 2D-E), suggesting that purine synthesis could be affected. SAM, which requires the purine nucleotide ATP for its synthesis, is also depleted^34^, and defects in purine synthesis have previously been shown to decrease SAM levels^35^. Indeed, levels of adenosine and guanosine nucleotides are decreased in MTR knockout cells (Figure 4A-B, Supplemental Figure 4A-B). Adenosine and guanosine nucleotide levels are further decreased in MTR knockout cells cultured in 5-methyl THF relative to those grown in folic acid, correlating with the observed reduction in proliferation rate of MTR knockout cells in 5-methyl THF (Figure 4A-B, Supplemental Figure 4A-B). The decrease in these nucleotides in the MTR knockout cells grown in folic acid may be related to the finding that some THF produced from folic acid is trapped as 5-methyl THF (Supplementary Figure 2A).

**Figure 4.**
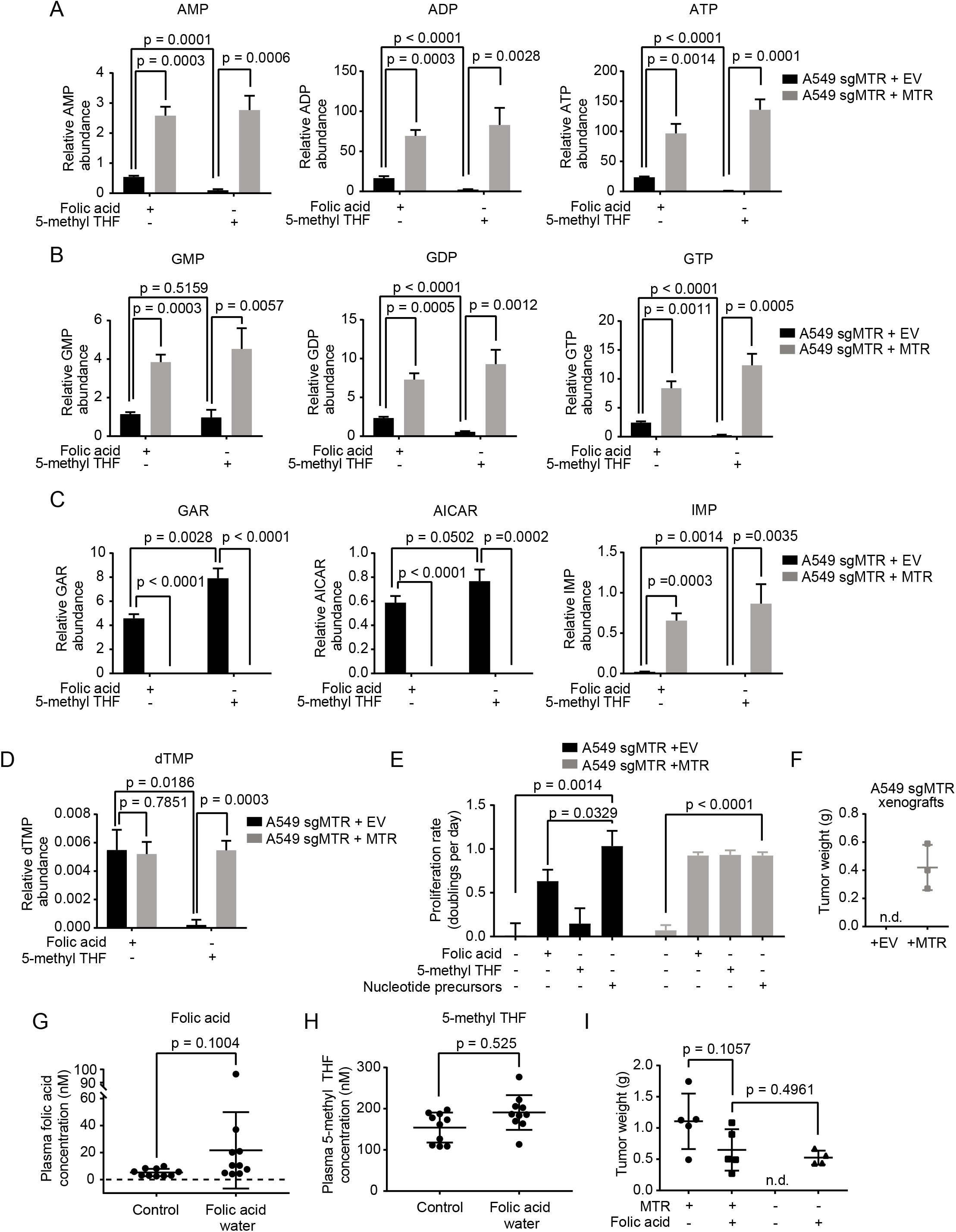
MTR is required to maintain nucleotide synthesis and for tumor formation. LC/MS measurement of intracellular AMP, ADP, ATP **(A)**, GMP, GDP, GTP **(B)**, GAR, AICAR, IMP **(C)**, and dTMP **(D)** levels in A549 cells without (+EV) or with MTR expression (+MTR) cultured for 4 days in the indicated folate. **(E)** Proliferation rates of A549 cells without (+EV) or with MTR expression (+MTR) in the indicated folate with or without the addition of 100 μM thymidine, 100 μM uridine, and 100 μM hypoxanthine (Nucleotide precursors). Data for the control conditions −FA, +FA, and 5-methyl THF are the same as those shown in Figure 3E. **(F)** Tumor weight of subcutaneous xenografts generated by injection of 100,000 A549 cells with no MTR expression (+EV) or expression of sgRNA resistant MTR (+MTR) into the flanks of NSG mice. Tumors were harvested 4 months after injection. n.d.: tumors were not detected. Plasma concentration of folic acid **(G)** and 5-methyl THF **(H)** in NSG mice provided with either normal water (Control) or water containing 0.1 g/L folic acid (Folic acid water) for 3 weeks. **(I)** Tumor weight of subcutaneous xenografts formed by injecting 1,000,000 A549 cells without or with MTR expression into the flanks of NSG mice provided with control water or water containing 0.1 g/L folic acid, beginning on the day the cells were injected. Tumors were harvested 3 months after injection. n.d.: tumors were not detected. Mean +/- SD is displayed for all panels. p values indicated on all panels are derived from two-tailed, unpaired Welch’s t tests. For all LC/MS measurements, data are normalized to the total protein content of cells in each condition and to an internal standard.

If nucleotide synthesis is impaired due to a lack of folate species derived from THF, intermediates prior to steps in nucleotide synthesis that utilize folates are predicted to accumulate (Supplemental Figure 4C)^36^. Consistent with this hypothesis, levels of two intermediates in purine synthesis, glycinamide ribonucleotide (GAR) and 5-amino-4-imidazolecarboxamide ribonucleotide (AICAR), are elevated in MTR knockout cells (Figure 4C, Supplemental Figure 4D). GAR and AICAR levels further increase in MTR knockout cells cultured in 5-methyl THF relative to cells grown in folic acid, consistent with the possibility that purine synthesis in these cells is limited by folate availability. Further supporting this hypothesis, inosine monophosphate (IMP), a purine precursor downstream of GAR and AICAR, is depleted in MTR knockout cells, particularly when grown in 5-methyl THF (Figure 4C, Supplemental Figure 4D).

Synthesis of the pyrimidine nucleotide deoxythymidine monophosphate (dTMP) requires the folate species 5,10-methylene THF (Figure 1A), which is unstable and not measurable directly by LC/MS without further derivatization^32^. If folate metabolism is globally disrupted by MTR knockout, cells would be predicted to also have low levels of 5,10-methylene THF and therefore exhibit impaired pyrimidine synthesis. Indeed, dTMP levels are depleted in MTR knockout cells grown in 5-methyl THF (Figure 4D, Supplemental Figure 4E). Strikingly, addition of the exogenous nucleotide precursors uridine, thymidine, and hypoxanthine rescues proliferation of MTR knockout cells in the absence of any folate source, suggesting that decreased MTR knockout cell proliferation is caused by nucleotide depletion (Figure 4E, Supplemental Figure 4F).

5-methyl THF is the predominant folate in circulation in humans^4^ and mice (Figure 1B); however, folic acid is also present at low levels in the circulation of both humans^4^ and mice (Figure 1B), and it is unknown whether circulating folic acid is sufficient to sustain tumor growth in the absence of MTR. To examine whether MTR is required for tumor growth given available folates *in vivo*, A549 cells with or without MTR expression were implanted into the flanks of immunocompromised NOD.Cg-*Prkdc^scid^Il2rg^tm1Wjl^*/SzJ (NSG) mice that were fed a diet containing folic acid at levels consistent with a typical human diet^37,38^. MTR expression was required for tumor formation in these conditions (Figure 4F). To test whether MTR is essential for tumor growth due to its role in utilizing circulating 5-methyl THF, mice were supplemented with excess folic acid, as this might provide a secondary folate source that circumvents the need for MTR-mediated conversion of 5-methyl THF to THF. Provision of folic acid in the drinking water of mice results in increased plasma folic acid levels in some mice (Figure 4G), without altering 5-methyl THF levels in circulation (Figure 4H). The degree of increase in plasma folic acid levels was variable and did not result in a statistically significant increase in average concentration (Figure 4G); however, folic acid supplementation was sufficient to rescue growth of MTR knockout tumors (Figure 4I), arguing that MTR is essential for tumor growth *in vivo* due to its role in producing THF from 5-methyl THF.

## DISCUSSION

Most commercial culture media formulations contain folic acid as the folate source. Though folic acid is capable of supporting rapid cell proliferation, it diminishes reliance on MTR, highlighting how the folate source in culture can alter dependence on folate-related enzymes and could impact the action of anti-folate drugs. For instance, inhibition of dihydrofolate reductase (DHFR) may be more toxic to cells cultured in folic acid, as conversion of folic acid to THF requires DHFR activity. This may explain the observation of increased sensitivity to DHFR inhibitors in culture compared to what is found in humans^39^.

Dietary methionine limitation restricts tumor growth and synergizes with anti-folate therapies in some cancers^40,41^. Methionine is an essential amino acid for mammals, but can be resynthesized from precursors involved in one-carbon and polyamine metabolism^42^. On a whole-animal level, MTR contributes to methionine re-synthesis; however, animals are capable of maintaining methionine levels via other pathways in the absence of MTR^9^, and we observe that MTR is not critical for maintaining intracellular methionine levels in a cancer cell-autonomous fashion. Further, despite the presence of sufficient methionine, SAM and SAH levels are depleted in MTR knockout cells. This is likely because decreased ATP availability can lower SAM levels, as has been observed previously^35^. Interestingly, we observe that significant ATP and SAM depletions are compatible with proliferation, suggesting that these metabolites may be present in excess in cultured cells.

MTR expression is essential for cancer cells to form tumors, suggesting that inhibition of MTR could be used as cancer therapy. Effective targeting of MTR would require tumor cells to have a higher demand for MTR activity than other tissues. The success of other therapies that influence folate metabolism^43–46^ suggests that this may be the case, and tumor types with a high demand for nucleotide synthesis might be expected to be most sensitive to MTR inhibition. Notably, MTR inhibition may not have the same effect as other anti-folates. For instance, non-squamous histologies of non-small cell lung cancer (NSCLC) are treated with the anti-folate pemetrexed^47^, and increased thymidylate synthase expression in squamous NSCLC^48,49^ correlates with reduced sensitivity to pemetrexed ^50^. However, squamous NSCLC cells do not similarly exhibit MTR overexpression^51^, possibly due to the complexities of MTR synthesis and cofactor regulation^52,53^; MTR function is supported by multiple enzymes that process and transport cobalamin, as well as a protein that specifically chaperones^54^ and regenerates MTR^55^. The use of models with physiological folates will be critical to define those tumor types that are most likely to respond to MTR inhibition.

## ACKNOWLEDGEMENTS

We thank Naama Kanarek for her comments and technical advice on folate measurements. We acknowledge all of the members of the Vander Heiden lab for input and advice on the manuscript. M.R.S. was supported by T32-GM007287 and by an MIT Koch Institute Graduate Fellowship. A.M.D. was supported by a Jane Coffin Childs Postdoctoral Fellowship. M.G.V.H. acknowledges support from R35CA242379, R01CA201276, the Ludwig Center at MIT, the MIT Center for Precision Cancer Medicine, SU2C, the Emerald Foundation, the Lustgarten Foundation, and a Faculty Scholars Grant from HHMI.

## AUTHOR CONTRIBUTIONS

Conceptualization, M.R.S., M.G.V.H.; Methodology, M.R.S., M.F.R., C.A.L.; Formal Analysis, M.R.S., A.M.D.; Investigation, M.R.S., A.M.D., M.F.R.; Resources, C.A.L.; Visualization, M.R.S., A.M.D.; Writing – Original Draft, M.R.S.; Writing – Review & Editing, M.R.S., A.M.D., M.G.V.H.; Funding Acquisition, M.R.S., A.M.D., M.G.V.H.

## DECLARATION OF INTERESTS

M.G.V.H. is on the scientific advisory board of Agios Pharmaceuticals, Aeglea Biotherapeutics, iTeos Therapeutics, and Auron Therapeutics. The other authors declare no competing interests.

All data generated or analyzed in this study are included in this article.

## MATERIALS AND METHODS

### Cell culture

Cells were passaged in RPMI-1640 (Corning Life Sciences, Tewksbury, MA) with 10% fetal bovine serum (FBS) that had been heat inactivated for 30 min. at 56°C (VWR Seradigm, Lot 120B14). All cells were cultured in a Heracell (Thermofisher) humidified incubators at 37°C and 5% CO_2_. A549 cells were obtained from ATCC (Manassas, VA). A549 cells are derived from a lung adenocarcinoma in a male patient. T.T cells were derived from an oral metastasis of an esophageal squamous cell carcinoma in a male patient. All cell lines were regularly tested for mycoplasma contamination using the Mycoprobe mycoplasma detection kit (R and D Systems, Minneapolis, MN). For experiments, cells were grown in RPMI-1640 without phenol red with 10% FBS that has been dialyzed to remove small molecules (Gibco, 26400044; batch 1: Lot #1841165, batch 2: Lot #2093857). We noted that different lots of dialyzed serum yielded subtle differences in sensitivity to folate starvation, so comparisons were only made between experiments carried out using the same lot of dialyzed serum. RPMI-1640 lacking folic acid was made using the method outlined in^56^. Briefly, enough of all of the components of RPMI-1640 media except for folic acid were weighed out to make 25 L of media, then the resulting powder was homogenized using an electric blade coffee grinder (Hamilton Beach, Glen Allen, VA, 80365) that had been washed with methanol then water. The resulting powder was resuspended in water to make RPMI-1640 media lacking folic acid and the pH was adjusted to ~7.6 - 7.8 with 1M HCl if necessary to improve solubilization. Folic acid (Sigma-Aldrich, St. Louis, MO, F8758) or (6S)-5-Methyl-5,6,7,8-tetrahydrofolic acid (5-methyl THF) (Schircks Laboratories, Bauma, Switzerland, 16.236) was dissolved in water to make 44-fold concentrated 100 uM stock solutions, then added back to RPMI-1640 media lacking folic acid to yield a final concentration of 2.26 μM, which is the standard concentration of folic acid in RPMI-1640. For experiments in which excess methionine was added to media, a 100 mM stock of methionine (Sigma-Aldrich, M5308) was generated in water and added back to RPMI-1640 media at 1 mM with the indicated folate. SAM (Sigma-Aldrich, A7007) was dissolved directly in cell culture media to 1 mM and sterile filtered. Stocks of the nucleosides uridine (Sigma-Aldrich, U3003) and thymidine (Sigma-Aldrich, T1895) were made at 100 mM (1000-fold concentrated) in water; hypoxanthine (Sigma-Aldrich, H9636) was dissolved at 20 mM (200-fold concentrated) in 0.1 N NaOH in water.

### Proliferation rates

Cellular proliferation rate in different media conditions was determined as previously described^57^. Cell lines were pre-starved of folic acid for 3 days, a time period that was empirically determined to be the longest folic acid starvation period that still allowed for full recovery of maximal proliferation rate when cells were provided a folate source. Note that cell lines lacking MTR expression were not pre-starved as any period of folic acid starvation affected their growth rate and recovery. For the 3-day pre-starvation, cells were plated into RPMI-1640 lacking folic acid at a density of 300,000 cells per 10 mL such that the cells were proliferating in log phase after 3 days. Cells were trypsinized, counted and plated into six well dishes (Corning Life Sciences) in 2 mL of RPMI-1640 medium lacking folic acid and incubated overnight. Initial seeding density was 40,000-50,000 cells/well. The next day, a six well plate of cells was trypsinized and counted to provide a number of cells at the start of the experiment. Cells were then washed twice with 2 mL of phosphate buffered saline (PBS), and 2 mL of the indicated media was added. For rescue experiments, 1 mM methionine, 1 mM SAM (Sigma-Aldrich, A7007), or the combination of 100 μM each uridine, thymidine, and hypoxanthine were added to the experimental media. Uridine, thymidine, and hypoxanthine were chosen for nucleotide precursor rescue experiments because they are readily taken up and utilized by cells, and can replace all *de novo* nucleotide synthesis pathways that require folates. Cells were then trypsinized and counted 4 days after adding the indicated media. Proliferation rate was determined using the following formula: Proliferation rate in doublings/day = [Log2(Final Day 4 cell count/Initial Day 0 cell count)]/4 days. Cells were counted using a Cellometer Auto T4 Plus Cell Counter (Nexcelom Bioscience, Lawrence, MA). All proliferation experiments were carried out in biological triplicate unless otherwise indicated on figure legends.

### Generation of MTR knockout cells

MTR knockout using CRISPR-Cas9 was accomplished using the pLenti-CRISPR v2 plasmid (Addgene Plasmid 49535)^58^. sgRNAs were designed based on previously described algorithms^59^, and MTR knockout was carried out using the following sgRNA sequence: TGGCATTGATCTCATCCCGC. Cell lines were diluted in 96 well plates to obtain single cell clones, and loss of MTR expression in individual clones was confirmed by western blot. sgRNA resistant cDNA of MTR was ordered from VectorBuilder (Shenandoah, TX) with silent mutations in the region of the gene targeted by the sgRNA.

### Western blot analysis

Cells were scraped in 300 μL - 1 mL RIPA buffer [25 mM Tris-Cl, 150 mM NaCl, 0.5% sodium deoxycholate, 1% Triton X-100, 1x cOmplete protease inhibitor (Roche, Basel, Switzerland), 1x phosSTOP (Sigma-Aldrich)]. The resulting lysate was clarified by centrifugation at 21000 x g for 10 min. Protein concentration of the lysate was determined by BCA assay (Thermofisher). Lysates were resuspended in Laemmli SDS-PAGE sample loading buffer (10% glycerol, 2% SDS, 60 mM Tris-Cl pH 6.8, 1% b-mercaptoethanol, 0.01% bromophenol blue) and denatured at 100°C for 5-10 min. Extracts were resolved by SDS-PAGE using 12% acrylamide gels running at 120 V until the dye front left the gel. After SDS-PAGE resolution, protein extracts were transferred to nitrocellulose using an iBlot semi-dry transfer system (Thermofisher; for MTR expression Western blots) or a Trans-Blot^®^ SD Semi-Dry Electrophoretic Transfer Cell (Bio-Rad; for mTOR signaling Western blots). Membranes were blocked in 5% non-fat dry milk or 5% BSA for phospho-epitope antibodies, incubated in primary antibodies to MTR (Abcam, Cambridge, UK, ab66039, 1:1000), β-actin (Cell Signaling Technology, Danvers, MA, 8457, 1:1000), phospho-T389 S6K (Cell Signaling Technology, Danvers, MA, 9234, 1:1000) S6K (Cell Signaling Technology, Danvers, MA, 9202, 1:1000), phospho-S235/6 RPS6 (Cell Signaling Technology, Danvers, MA, 4858, 1:1000), RPS6 (Cell Signaling Technology, Danvers, MA, 2217, 1:1000), or mono/dimethyl lysine residues (Abcam, Cambridge UK, ab23366, 1:1000) overnight and detected using HRP-conjugated secondary antibodies and chemiluminescence with the Western Lightning Plus-ECL Reagent (Perkin Elmer).

### Metabolite extraction

For analysis of mouse plasma folates, 10 μL of plasma was mixed with 90 μL extraction buffer (80:20 methanol:water with 2.5 mM sodium ascorbate, 25 mM ammonium acetate, 100 nM aminopterin). Samples were vortexed for 10 minutes at 4 °C, then centrifuged at 21000 x g at 4°C for 10 minutes. Supernatant was removed and dried under nitrogen. For extraction of polar metabolites, 5 μL of a standard composed of a ^13^C-labeled amino acid mix (Cambridge Isotopes, Tewksbury, MA, MSK-A2-1.2) diluted to a concentration of 200 μM per amino acid was added to 600 μL 80% HPLC grade methanol (Sigma-Aldrich, 646377-4X4L) and added to cells after draining media and washing with 2-3 mL ice cold saline. Cells were then scraped, transferred to an Eppendorf tube, vortexed for 10 minutes at 4 °C, then centrifuged at 21000 x g at 4°C for 10 minutes. 400 μL of sample was removed and dried under nitrogen. Extraction of intracellular folates was performed according to previously described methods^60^. 1 mL 80% HPLC grade methanol with 2.5 mM sodium ascorbate and 25 mM ammonium acetate, with a 100 nM aminopterin standard and 500 nM each of a ^13^C-labeled amino acid mix, was added to cells in a 6 cm dish on ice after draining media and washing with 5 mL ice cold saline. Cells were then scraped, transferred to an Eppendorf tube, vortexed briefly, and centrifuged at 21000 x g at 4°C for 10 minutes. 600 μL of sample was removed for analysis of folates and 100 uL was removed for analysis of polar metabolites; both were dried under nitrogen. Folate samples were resuspended in 500 uL rat serum (Sigma-Aldrich, R9759) for 2 hours at 37°C that had been stripped twice with 0.5 g activated charcoal (Sigma-Aldrich, C9157) per 10 mL serum, centrifuged at 4000 RPM at 4°C for 10 minutes to sediment charcoal and filtered each time after stripping through a 0.45 uM PES filter. 5 uL 1:1 formic acid to water was added to folate samples to adjust pH, and samples were cleaned up on SPE columns (Agilent Technologies, 14102062) according to the manufacturer’s instructions and dried under nitrogen.

### LC/MS analysis of non-folate metabolites

Dried samples were resuspended in 100 μL HPLC grade water. LC-MS analysis was performed using a QExactive orbitrap mass spectrometer using an Ion Max source and heated electro-spray ionization (HESI) probe coupled to a Dionex Ultimate 3000 UPLC system (Thermofisher). External mass calibration was performed every 7 days. Polar metabolite samples were separated by chromatography by injecting 10 μL of sample on a SeQuant ZIC-pHILIC 2.1 mm x 150 mm (5 μm particle size) column. Flow rate was set to 150 mL/min. and temperatures were set to 25°C for the column compartment and 4°C for the autosampler tray. Mobile phase A was 20 mM ammonium carbonate, 0.1% ammonium hydroxide. Mobile phase B was 100% acetonitrile. The chromatographic gradient was: 0–20 min.: linear gradient from 80% to 20% mobile phase B; 20–20.5 min.: linear gradient from 20% to 80% mobile phase B; 20.5 to 28 min.: hold at 80% mobile phase B. The mass spectrometer was operated in full scan, polarity-switching mode and the spray voltage was set to 3.0 kV, the heated capillary held at 275°C, and the HESI probe was held at 350°C. The sheath gas flow rate was 40 units, the auxiliary gas flow was 15 units and the sweep gas flow was one unit. The MS data acquisition was performed in a range of 70–1000 m/z, with the resolution set set at 70,000, the AGC target at 1×10^6^, and the maximum injection time at 20 msec. Relative quantitation of metabolites was performed with XCalibur QuanBrowser 2.2 (Thermo Fisher Scientific) using a 5 ppm mass tolerance and referencing an in-house library of chemical standards. Peak areas were normalized to cell number and ^13^C-amino acid standard peak areas. All LC/MS measurements of non-folate metabolites were carried out in biological triplicate, with the exception of Supplemental Figure 1D-F, for which n=1 at each time point.

### LC/MS analysis of folate species

Detection of folate species was performed on the same instrumentation described above, as outlined in previous work^60^. LC-MS measurements were initiated on the same day as extraction. In general, instrument settings remained the same unless specified. Samples were resuspended in 50 μL water and 15 μL was injected onto an Ascentis^®^ Express C18 HPLC column (2.7 μm × 15 cm × 2.1 mm; Sigma Aldrich). The column oven and autosampler tray were held at 30°C and 4°C, respectively. The following conditions were used to achieve chromatographic separation: Buffer A was 0.1% formic acid; buffer B was acetonitrile with 0.1% formic acid. The chromatographic gradient was run at a flow rate of 0.250 mL/min as follows: 0-5min.: gradient was held at 5% B; 5-10 min.: linear gradient of 5% to 36% B; 10.1-14.0 min.: linear gradient from 36%-95% B; 14.1-18.0 min.: gradient was returned to 5% B. The mass spectrometer was operated in full-scan, positive ionization mode. MS data acquisition was performed using three narrow-range scans: 438-450 m/z; 452-462 m/z; and 470-478 m/z, with the resolution set at 70,000, the AGC target at 10e6, and the maximum injection time of 150 msec. Relative quantitation of folate species was performed with XCalibur QuanBrowser 2.2 (Thermo Fisher Scientific) using a 5 ppm mass tolerance. Folate species were identified using chemical standards, and 5,10-methenyl THF and 10-formyl THF were pooled during analysis due to the possibility of interconversion of the species in each pool after lysis^61^. LC/MS measurements of folates were carried out in biological triplicate, with the exception of Figure 1D and Supplemental Figure 2A, for which n=1 at each time point.

### Mouse tumor studies

Cells were trypsinized and either 100,000 or 1,000,000 cells were resuspended in 100 μL PBS as indicated on figure legends. Cells were injected into the flanks of NSG mice. Mice were euthanized according to institutional guidelines and tumor weight was recorded. 0.1 g/L folic acid water was prepared by resuspending folic acid in tap water, then filtering the mixture through a 0.22 μm filter to sterilize the water. All animal studies were carried out according to MIT Committee on Animal Care guidelines.

### Blood collection from mice

Blood was collected from fed, anesthetized mice by retro-orbital bleeding at 11 AM. Blood was placed directly into EDTA coated collection tubes (Sarstedt, Nümbrecht, Germany, 41.1395.105) and centrifuged 10’ at 845 x g; the supernatant of plasma was transferred to another tube.

## SUPPLEMENTAL FIGURE LEGENDS

**Supplemental Figure 1.**
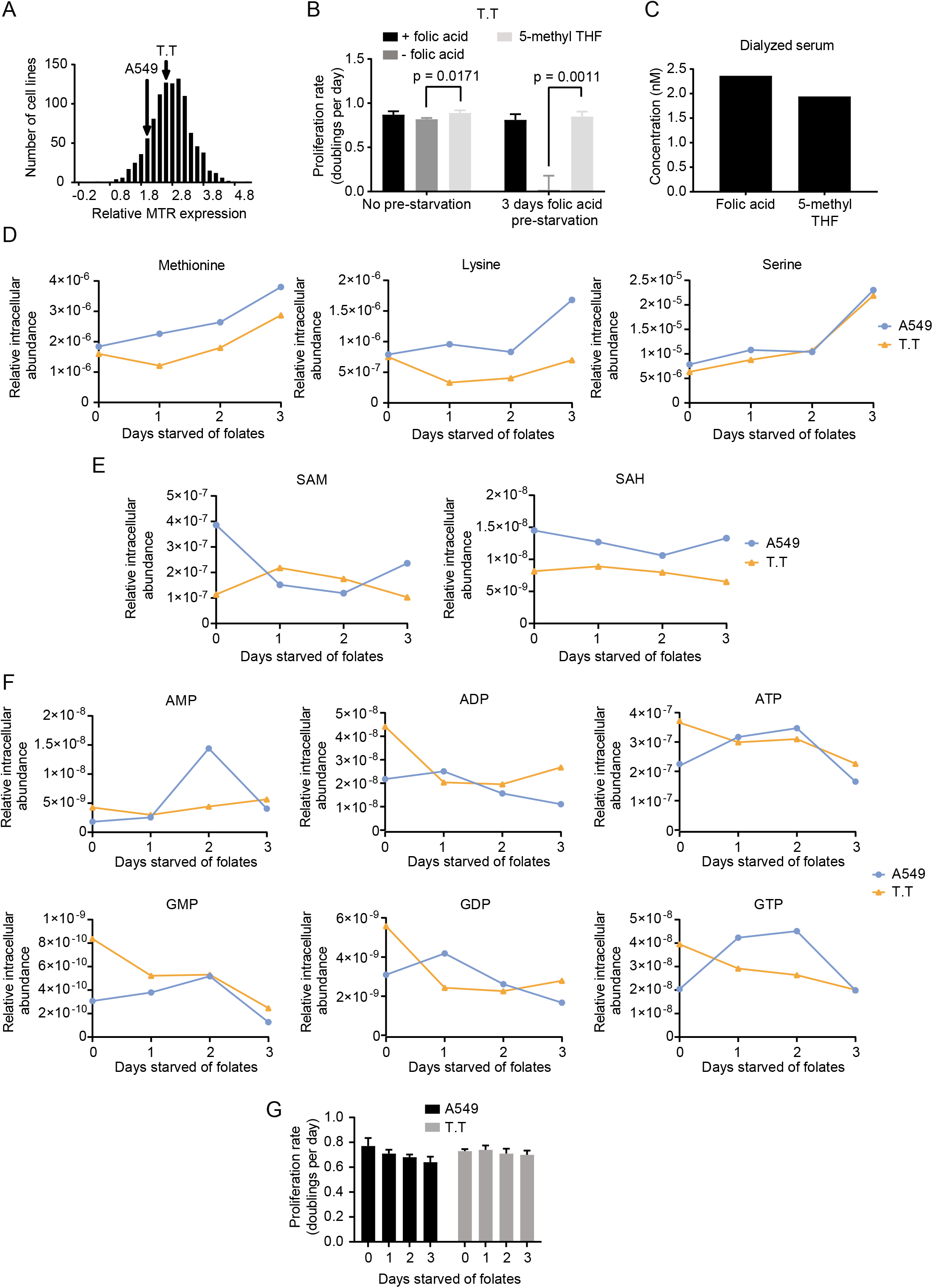
Characterization of cells cultured in 5-methyl THF. **(A)** Histogram of MTR mRNA expression levels in cell lines as reported in The Cancer Genome Atlas^23^. **(B)** Proliferation rate of T.T cells in media with the indicated folate before (no pre-starvation) or after 3 days of culture with no folic acid (pre-starvation). **(C)** LC/MS measurement of folic acid and 5-methyl THF concentrations present in the dialyzed fetal bovine serum used to prepare culture medium for this study (batch 1; see Methods). **(D)** LC/MS measurement of intracellular methionine, lysine, and serine levels in A549 and T.T cells cultured for the indicated number of days in media lacking any added folates. LC/MS measurement of intracellular SAM and SAH levels **(E)** and nucleotide levels **(F)** in A549 and T.T cells cultured for the indicated number of days in media lacking any added folates. **(G)** Proliferation rate of A549 or T.T cells cultured in media lacking any added folates for the indicated number of days. Mean +/- SD is displayed for all panels. p values indicated on all panels are derived from two-tailed, unpaired Welch’s t tests. For all LC/MS measurements, data are normalized to cell number and to an internal standard.

**Supplemental Figure 2.**
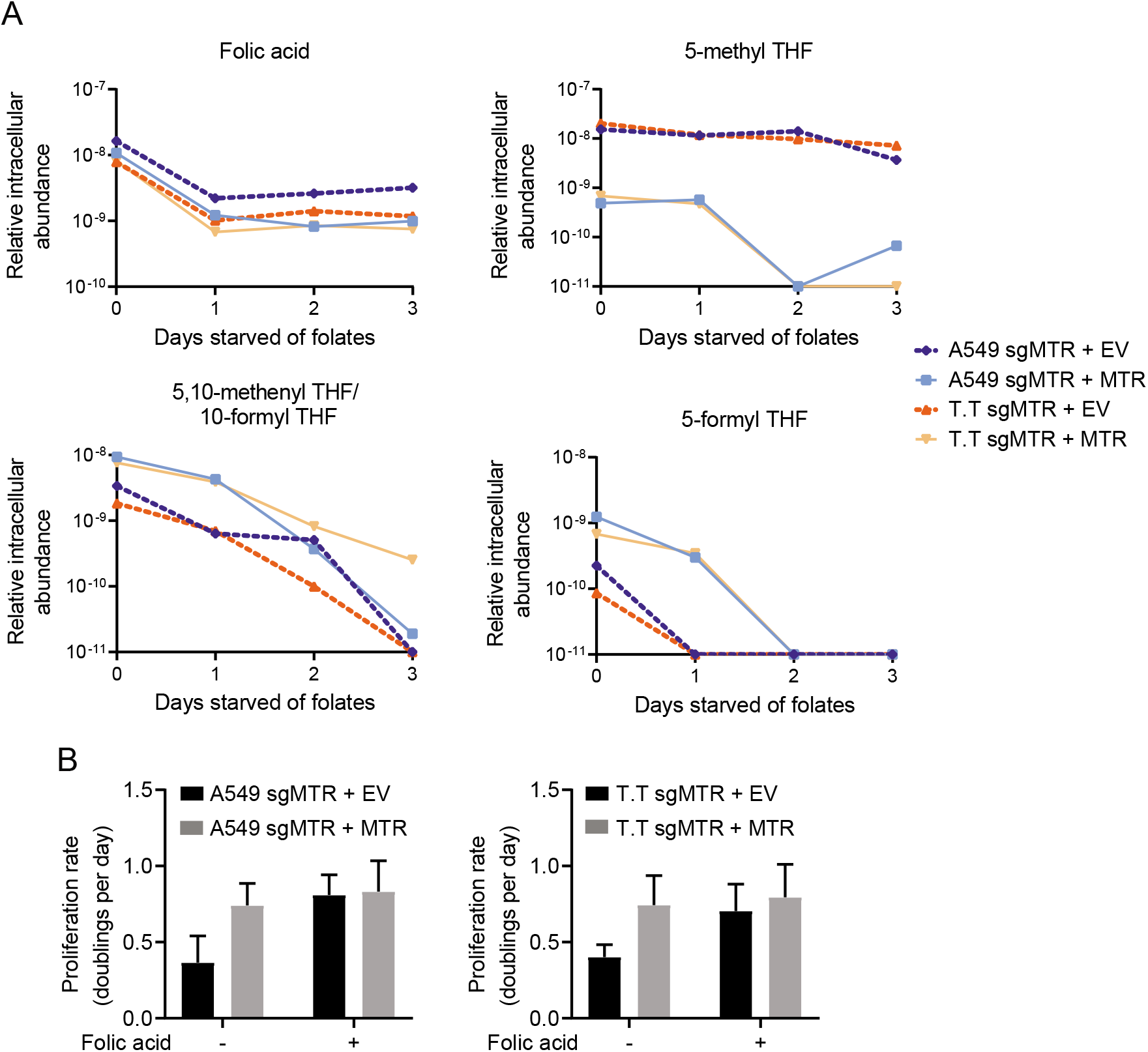
MTR expression affects levels of intracellular folate species. **(A)** LC/MS measurement of intracellular folic acid, 5-methyl THF, 5,10-methenyl THF/10-formyl THF, and 5-formyl THF levels in A549 and T.T cells with no MTR expression (+EV) or expression of sgRNA resistant MTR (+MTR) cultured for the indicated number of days in media lacking any added folates. Data are normalized to the total cell number in each condition and to an internal standard. **(B)** Proliferation rate of A549 or T.T cells without (+EV) or with MTR expression (+MTR) cultured in media lacking any added folates for 3 days. Mean +/- SD is displayed.

**Supplemental Figure 3.**
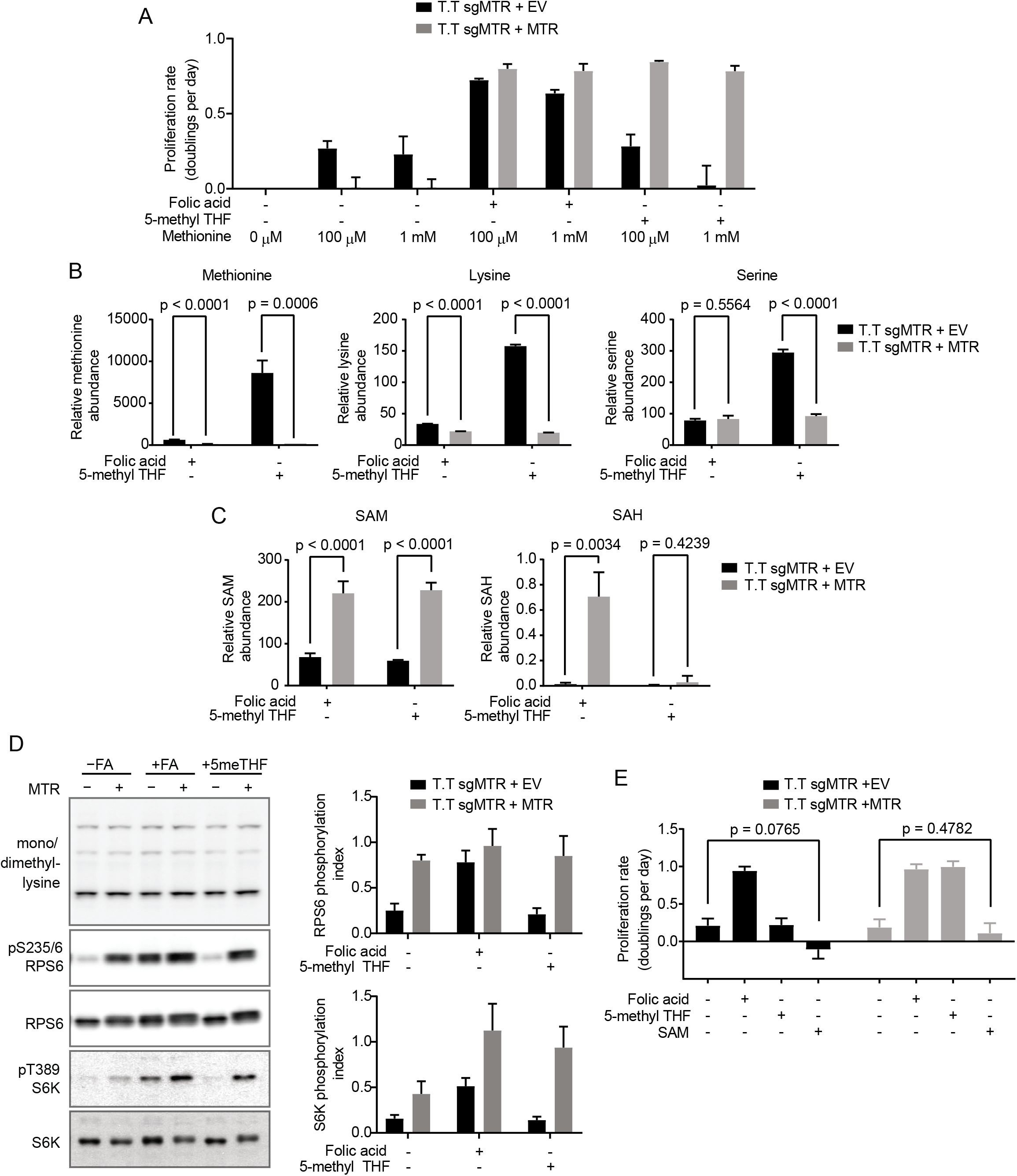
MTR expression affects SAM but not methionine metabolism in T.T cells. **(A)** Proliferation rates of T.T cells without (+EV) or with MTR expression (+MTR) cultured in the indicated folate with or without the addition of 1 mM methionine to supplement the 100 μM methionine present in RPMI-1640 culture media. **(B)** LC/MS measurement of intracellular methionine, lysine, and serine levels in T.T cells without (+EV) or with MTR expression (+MTR) cultured for 4 days in the indicated folate. **(C)** LC/MS measurement of intracellular SAM and SAH levels in T.T cells without (+EV) or with MTR expression (+MTR) cultured for 4 days in the indicated folate. **(D)** Western blots to assess levels of mono- or dimethyl lysine-containing proteins, RPS6 phosphorylated at S235/6, total RPS6, S6K phosphorylated at T389, and total S6K in T.T cells without (+EV) or with MTR expression (+MTR). Phosphorylation index is defined as the phosphorylated protein antibody signal divided by the same total protein antibody signal in each lane. **(E)** Proliferation rates of T.T cells without (+EV) or with MTR expression (+MTR) cultured in the indicated folate with or without the addition of 1 mM SAM. Mean +/- SD is displayed for all panels. p values indicated on all panels are derived from two-tailed, unpaired Welch’s t tests. For LC/MS measurements, data are normalized to the total protein content of cells in each condition and to an internal standard.

**Supplemental Figure 4.**
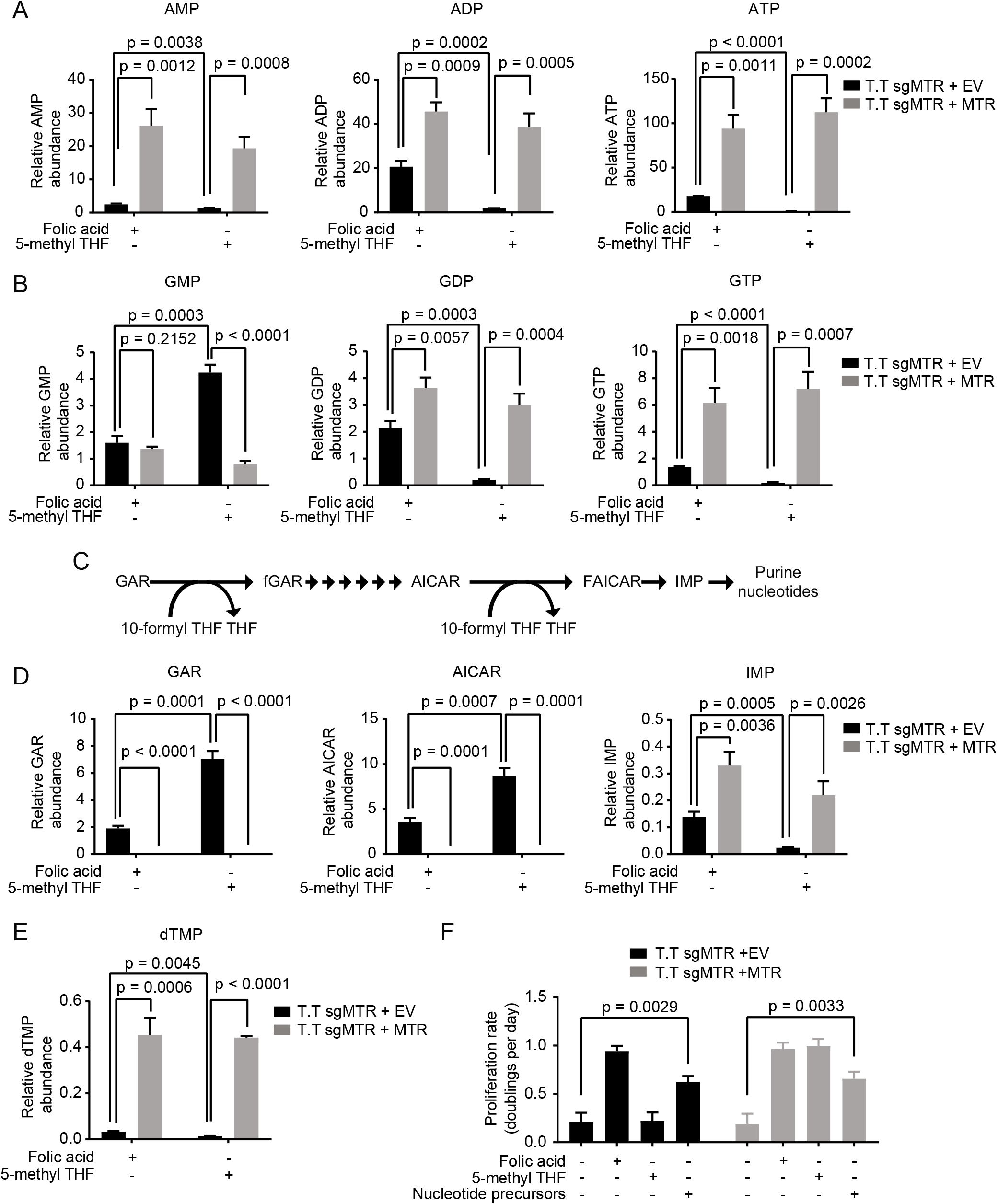
MTR knockout results in folate insufficiency and impairs nucleotide synthesis in T.T cells. LC/MS measurement of intracellular AMP, ADP, ATP **(A)**, GMP, GDP, and GTP **(B)**, levels in T.T cells without (+EV) or with MTR expression (+MTR) cultured for 4 days in the indicated folate. **(C)** Schematic showing select reactions involved in purine synthesis. GAR: glycinamide ribonucleotide. fGAR: 5’-phosphoribosyl-N-formylglycinamide. AICAR: 5-amino-4-imidazolecarboxamide ribonucleotide. FAICAR: 5-formamidoimidazole-4-carboxamide ribonucleotide. IMP: inosine monophosphate. THF: tetrahydrofolate. LC/MS measurement of intracellular GAR, AICAR, IMP **(D)**, and dTMP **(E)** levels in T.T cells without (+EV) or with MTR expression (+MTR) cultured for 4 days in the indicated folate. **(F)** Proliferation rates of T.T cells without (+EV) or with MTR expression (+MTR) cultured in the indicated folate with or without the addition of 100 μM thymidine, 100 μM uridine, and 100 μM hypoxanthine (Nucleotide precursors). Data for the control conditions – FA, +FA, and 5-methyl THF are the same as those shown in Supplemental Figure 3E. Mean +/- SD is displayed for all panels. p values indicated on all panels are derived from two-tailed, unpaired Welch’s t tests. For all LC/MS measurements, data are normalized to the total protein content of cells in each condition and to an internal standard.

## REFERENCES

1 Chiang, P. K. et al. S-Adenosylmethionine and methylation. FASEB J 10, 471–480 (1996).

2 Lane, A. N. & Fan, T. W. Regulation of mammalian nucleotide metabolism and biosynthesis. Nucleic Acids Res 43, 2466–2485, doi:10.1093/nar/gkv047 (2015).

3 Ducker, G. S. & Rabinowitz, J. D. One-Carbon Metabolism in Health and Disease. Cell Metab 25, 27–42, doi:10.1016/j.cmet.2016.08.009 (2017).

4 Pfeiffer, C. M. et al. Folate status and concentrations of serum folate forms in the US population: National Health and Nutrition Examination Survey 2011-2. Br J Nutr 113, 1965–1977, doi:10.1017/S0007114515001142 (2015).

5 Muir, A., Danai, L. V. & Vander Heiden, M. G. Microenvironmental regulation of cancer cell metabolism: implications for experimental design and translational studies. Dis Model Mech 11, doi:10.1242/dmm.035758 (2018).

6 Rancati, G., Moffat, J., Typas, A. & Pavelka, N. Emerging and evolving concepts in gene essentiality. Nat Rev Genet 19, 34–49, doi:10.1038/nrg.2017.74 (2018).

7 Banerjee, R. V. & Matthews, R. G. Cobalamin-dependent methionine synthase. FASEB J 4, 1450–1459 (1990).

8 Stover, P. J. Vitamin B12 and older adults. Curr Opin Clin Nutr Metab Care 13, 24–27, doi:10.1097/MCO.0b013e328333d157 (2010).

9 Chanarin, I., Deacon, R., Lumb, M., Muir, M. & Perry, J. Cobalamin-folate interrelations: a critical review. Blood 66, 479–489 (1985).

10 Fujii, K., Nagasaki, T. & Huennekens, F. M. Accumulation of 5-methyltetrahydrofolate in cobalamin-deficient L1210 mouse leukemia cells. J Biol Chem 257, 2144–2146 (1982).

11 Walker, P. R. et al. Induction of apoptosis in neoplastic cells by depletion of vitamin B12. Cell Death Differ 4, 233–241, doi:10.1038/sj.cdd.4400225 (1997).

12 McLean, G. R. et al. Cobalamin analogues modulate the growth of leukemia cells in vitro. Cancer Res 57, 4015–4022 (1997).

13 Corcino, J. J., Zalusky, R., Greenberg, M. & Herbert, V. Coexistence of Pernicious Anaemia and Chronic Myeloid Leukaemia: An Experiment of Nature Involving Vitamin BI2 Metabolism. Br J Haematol 20, 511 (1971).

14 Eastwood, D. W., Green, C. D., Lambdin, M. A. & Gardner, R. Effect of Nitrous Oxide on the White-Cell Count in Leukemia. New England Journal of Medicine 268, 297–299, doi:10.1056/nejm196302072680607 (1963).

15 Ikeda, K. et al. Antileukemic effect of nitrous oxide in a patient with chronic myelogenous leukemia. Am J Hematol 30, 114 (1989).

16 Buckel, W. in Encyclopedia of Life Sciences (2007).

17 Matthews, J. H. Cyanocobalamin [c-lactam] Inhibits Vitamin B12 and Causes Cytotoxicity in HL60 Cells: Methionine Protects Cells Completely. Blood 89, 4600–4607 (1997).

18 Liteplo, R. G., Hipwell, S. E., Rosenblatt, D. S., Sillaots, S. & Lue-Shing, H. Changes in cobalamin metabolism are associated with the altered methionine auxotrophy of highly growth autonomous human melanoma cells. J Cell Physiol 149, 332–338, doi:10.1002/jcp.1041490222 (1991).

19 Boss, G. R. Cobalamin inactivation decreases purine and methionine synthesis in cultured lymphoblasts. J Clin Invest 76, 213–218, doi:10.1172/JCI111948 (1985).

20 Lumb, M. et al. Effects of nitrous oxide-induced inactivation of cobalamin on methionine and S-adenosylmethionine metabolism in the rat. Biochim Biophys Acta 756, 354–359 (1983).

21 van der Westhuyzen, J., Fernandes-Costa, F. & Metz, J. Cobalamin inactivation by nitrous oxide produces severe neurological impairment in fruit bats: protection by methionine and aggravation by folates. Life Sci 31, 2001–2010 (1982).

22 Scott, J. M., Dinn, J. J., Wilson, P. & Weir, D. G. Pathogenesis of subacute combined degeneration: a result of methyl group deficiency. Lancet 2, 334–337 (1981).

23 Cancer Genome Atlas Research, N. et al. The Cancer Genome Atlas Pan-Cancer analysis project. Nat Genet 45, 1113–1120, doi:10.1038/ng.2764 (2013).

24 Hansen, M. F., Jensen, S. O., Fuchtbauer, E. M. & Martensen, P. M. High folic acid diet enhances tumour growth in PyMT-induced breast cancer. Br J Cancer 116, 752–761, doi:10.1038/bjc.2017.11 (2017).

25 Farber, S. et al. The Action of Pteroylglutamic Conjugates on Man. Science 106, 619–621, doi:10.1126/science.106.2764.619 (1947).

26 Farber, S., Diamond, L. K., Mercer, R. D., Sylvester, R. F. & Wolff, J. A. Temporary remissions in acute leukemia in children produced by folic acid antagonist, 4-aminopteryl-glutamic acid (aminopterin). New England Journal of Medicine 238, 787–793 (1948).

27 Palmer, A. M., Kamynina, E., Field, M. S. & Stover, P. J. Folate rescues vitamin B12 depletion-induced inhibition of nuclear thymidylate biosynthesis and genome instability. Proc Natl Acad Sci U S A 114, E4095–E4102, doi:10.1073/pnas.1619582114 (2017).

28 Sullivan, M. R. et al. Increased Serine Synthesis Provides an Advantage for Tumors Arising in Tissues Where Serine Levels Are Limiting. Cell Metab 29, 1410–1421 e1414, doi:10.1016/j.cmet.2019.02.015 (2019).

29 Su, X., Wellen, K. E. & Rabinowitz, J. D. Metabolic control of methylation and acetylation. Curr Opin Chem Biol 30, 52–60, doi:10.1016/j.cbpa.2015.10.030 (2016).

30 Cho, H. S. et al. Enhanced HSP70 lysine methylation promotes proliferation of cancer cells through activation of Aurora kinase B. Nat Commun 3, 1072, doi:10.1038/ncomms2074 (2012).

31 Gu, X. et al. SAMTOR is an <em>S</em>-adenosylmethionine sensor for the mTORC1 pathway. Science 358, 813–818, doi:10.1126/science.aao3265 (2017).

32 Chen, L., Ducker, G. S., Lu, W., Teng, X. & Rabinowitz, J. D. An LC-MS chemical derivatization method for the measurement of five different one-carbon states of cellular tetrahydrofolate. Anal Bioanal Chem, doi:10.1007/s00216-017-0514-4 (2017).

33 Stover, P. & Schirch, V. Evidence for the accumulation of a stable intermediate in the nonenzymatic hydrolysis of 5,10-methenyltetrahydropteroylglutamate to 5-formyltetrahydropteroylglutamate. Biochemistry 31, 2148–2155, doi:10.1021/bi00122a036 (1992).

34 Mato, J. M., Alvarez, L., Ortiz, P. & Pajares, M. A. S-adenosylmethionine synthesis: molecular mechanisms and clinical implications. Pharmacol Ther 73, 265–280 (1997).

35 Maddocks, O. D., Labuschagne, C. F., Adams, P. D. & Vousden, K. H. Serine Metabolism Supports the Methionine Cycle and DNA/RNA Methylation through De Novo ATP Synthesis in Cancer Cells. Mol Cell 61, 210–221, doi:10.1016/j.molcel.2015.12.014 (2016).

36 Labuschagne, C. F., van den Broek, N. J., Mackay, G. M., Vousden, K. H. & Maddocks, O. D. K. Serine, but not glycine, supports one-carbon metabolism and proliferation of cancer cells. Cell Rep 7, 1248–1258, doi:10.1016/j.celrep.2014.04.045 (2014).

37 Reeves, P. G. Components of the AIN-93 Diets as Improvements in the AIN-76A Diet. J Nutr 127, 838–841 (1997).

38 Bailey, R. L. et al. Total folate and folic acid intake from foods and dietary supplements in the United States: 2003-2006. Am J Clin Nutr 91, 231–237, doi:10.3945/ajcn.2009.28427 (2010).

39 Gaukroger, J. et al. Paradoxical response of malignant melanoma to methotrexate in vivo and in vitro. Br J Cancer 47, 671–679, doi:10.1038/bjc.1983.105 (1983).

40 Chaturvedi, S., Hoffman, R. M. & Bertino, J. R. Exploiting methionine restriction for cancer treatment. Biochem Pharmacol 154, 170–173, doi:10.1016/j.bcp.2018.05.003 (2018).

41 Gao, X. et al. Dietary methionine influences therapy in mouse cancer models and alters human metabolism. Nature 572, 397–401, doi:10.1038/s41586-019-1437-3 (2019).

42 Sanderson, S. M., Gao, X., Dai, Z. & Locasale, J. W. Methionine metabolism in health and cancer: a nexus of diet and precision medicine. Nat Rev Cancer 19, 625–637, doi:10.1038/s41568-019-0187-8 (2019).

43 Sakura, T. et al. High-dose methotrexate therapy significantly improved survival of adult acute lymphoblastic leukemia: a phase III study by JALSG. Leukemia 32, 626–632, doi:10.1038/leu.2017.283 (2017).

44 Baldwin, C. M. & Perry, C. M. Pemetrexed: A Review in its Use in the Management of Advanced Non-Squamous Non-Small Cell Lung Cancer. Drugs 69, 2279–2302 (2009).

45 Gonen, N. & Assaraf, Y. G. Antifolates in cancer therapy: structure, activity and mechanisms of drug resistance. Drug Resist Updat 15, 183–210, doi:10.1016/j.drup.2012.07.002 (2012).

46 Visentin, M., Zhao, R. & Goldman, I. D. The antifolates. Hematol Oncol Clin North Am 26, 629–648, ix, doi:10.1016/j.hoc.2012.02.002 (2012).

47 Li, X., Wei, S. & Chen, J. Critical appraisal of pemetrexed in the treatment of NSCLC and metastatic pulmonary nodules. Onco Targets Ther 7, 937–945, doi:10.2147/OTT.S45148 (2014).

48 Otake, Y. et al. Expression of thymidylate synthase in human non-small cell lung cancer. Jpn J Cancer Res 90, 1248–1253 (1999).

49 Ceppi, P. et al. Squamous cell carcinoma of the lung compared with other histotypes shows higher messenger RNA and protein levels for thymidylate synthase. Cancer 107, 1589–1596, doi:10.1002/cncr.22208 (2006).

50 Takezawa, K. et al. Thymidylate synthase as a determinant of pemetrexed sensitivity in non-small cell lung cancer. Br J Cancer 104, 1594–1601, doi:10.1038/bjc.2011.129 (2011).

51 Li, Y. et al. Transcriptomic and functional network features of lung squamous cell carcinoma through integrative analysis of GEO and TCGA data. Sci Rep 8, 15834, doi:10.1038/s41598-018-34160-w (2018).

52 Gherasim, C., Lofgren, M. & Banerjee, R. Navigating the B(12) road: assimilation, delivery, and disorders of cobalamin. J Biol Chem 288, 13186–13193, doi:10.1074/jbc.R113.458810 (2013).

53 Oltean, S. & Banerjee, R. Nutritional modulation of gene expression and homocysteine utilization by vitamin B12. J Biol Chem 278, 20778–20784, doi:10.1074/jbc.M300845200 (2003).

54 Yamada, K., Gravel, R. A., Toraya, T. & Matthews, R. G. Human methionine synthase reductase is a molecular chaperone for human methionine synthase. Proc Natl Acad Sci U S A 103, 9476–9481 (2006).

55 Olteanu, H. & Banerjee, R. Human methionine synthase reductase, a soluble P-450 reductase-like dual flavoprotein, is sufficient for NADPH-dependent methionine synthase activation. J Biol Chem 276, 35558–35563, doi:10.1074/jbc.M103707200 (2001).

56 Muir, A. et al. Environmental cystine drives glutamine anaplerosis and sensitizes cancer cells to glutaminase inhibition. Elife 6, doi:10.7554/eLife.27713 (2017).

57 Sullivan, L. B. et al. Supporting Aspartate Biosynthesis Is an Essential Function of Respiration in Proliferating Cells. Cell 162, 552–563, doi:10.1016/j.cell.2015.07.017 (2015).

58 Sanjana, N. E., Shalem, O. & Zhang, F. Improved vectors and genome-wide libraries for CRISPR screening. Nat Methods 11, 783–784, doi:10.1038/nmeth.3047 (2014).

59 Hart, T. et al. High-Resolution CRISPR Screens Reveal Fitness Genes and Genotype-Specific Cancer Liabilities. Cell 163, 1515–1526, doi:10.1016/j.cell.2015.11.015 (2015).

60 Kanarek, N. et al. Histidine catabolism is a major determinant of methotrexate sensitivity. Nature 559, 632–636, doi:10.1038/s41586-018-0316-7 (2018).

61 Chen, L., Ducker, G. S., Lu, W., Teng, X. & Rabinowitz, J. D. An LC-MS chemical derivatization method for the measurement of five different one-carbon states of cellular tetrahydrofolate. Anal Bioanal Chem 409, 5955–5964, doi:10.1007/s00216-017-0514-4 (2017).

